# Altered visual responses in Drosophila BBSome mutants support a nonciliary role in the retina

**DOI:** 10.64898/2026.11.16.766922

**Authors:** Sarolta Szentes, Milena Damulewicz, Kinga Kapusi, Katalin Farkas, David M Glover, Levente Kovacs

## Abstract

The BBSome regulates ciliary membrane protein trafficking and mutations in its individual subunits lead in humans, to Bardet–Biedl syndrome, characterized by ciliopathic features including retinal degeneration. Here, we investigated BBSome function in *Drosophila melanogaster*, whose retinal photoreceptors lack cilia. We used CRISPR/Cas9 mutagenesis to generate null alleles of six of the conserved BBSome subunits and the associated regulatory molecule BBS3/ARL6. Each single mutant was viable and fertile and only the BBS1-BBS9 double mutant combination showed reduced male fertility. The BBSome mutants retained robust climbing ability and normal ciliary features in their sensory organs. However, all mutant genotypes showed reduced positive phototaxis. Locomotor recordings also demonstrated reduced light responsiveness in the mutants, with a transient suppression of activity during a light pulse in constant darkness. Fluorescently tagged wild-type BBSome subunits formed puncta in differentiating photoreceptors suggesting their association within the developing eye. In the adult mutants, a Rhodopsin 1-GFP reporter showed increased rhabdomeric fluorescence and additional puncta in the photoreceptor cell-body region which were not present in the wild-type. These findings support a non-ciliary contribution of BBSome proteins to orchestrating rhodopsin distribution and determining visual function. The altered localization of rhodopsin 1-GFP is consistent with disrupted rhodopsin trafficking or turnover, through a transport step independent of ciliary axonemes.

## Introduction

Bardet–Biedl syndrome is a categorized as a collection of congenital ciliopathies, characterized by retinal dystrophy, obesity, polydactyly and renal abnormalities, resulting from mutations in genes encoding proteins involved in ciliary function (Beales et al., 1999; Hsu et al., 2017). A conserved group of these proteins forms the BBSome, an eight-subunit complex comprising BBS1, BBS2, BBS4, BBS5, BBS7, BBS8, BBS9 and BBS18 (Nachury et al., 2007; Loktev et al., 2008). In addition, the small GTPase BBS3/ARL6 regulates recruitment of the complex to membranes rather than forming part of the BBsome’s structural core. Together, BBS3/ARL6 and the BBSome promote selective trafficking of membrane proteins, including the removal of activated receptors through the ciliary transition zone (Jin et al., 2010; Ye et al., 2018). Studies of vertebrate photoreceptors have established requirements for the BBSome in the formation and maintenance of the photosensitive outer segment, a specialized ciliary compartment. Loss of BBSome function also permits the accumulation of proteins normally excluded from this compartment, indicating a role in maintaining the compartment’s composition (Datta et al., 2015; Hsu et al., 2017). Experimental evidence also suggests that BBS proteins can also act outside cilia. Depletion of BBS1 reduces cell-surface leptin receptor levels in the mouse nervous system, whereas depletion of the intraflagellar transport protein IFT88 does not reproduce this effect (Guo et al., 2016). BBSome subunits also promote centrosome polarization during immune synapse assembly in nonciliated T cells (Cassioli et al., 2021). These findings extend the functions of BBSome proteins beyond the cilium, however, it remains unclear whether these non-ciliary functions are mediated by the BBSome complex as a whole or by individual BBSome proteins acting independently of the complex. In Drosophila, comparative genomic and expression analyses identified BBS1, BBS4 and BBS8 subunits in mechanosensory and chemosensory organs (Avidor-Reiss et al., 2004). However, Drosophila’s retinal photoreceptors lack cilia and concentrate rhodopsin in rhabdomere microvilli. Their photosensitive membranes nevertheless require regulated trafficking: Rab11 mediates post-Golgi rhodopsin delivery, whereas retromer-dependent recycling maintains rhodopsin homeostasis after endocytosis (Satoh et al., 2005; Wang et al., 2014). The fly therefore provides a means of examining BBSome-associated trafficking independently of a photoreceptor cilium. The above findings led us to ask whether the conserved BBSome machinery contributes to visual function in the nonciliated *Drosophila* retina. Here, we address this question by generating mutants for BBSome subunits and the associated regulator BBS3, and comparing sensory behaviour with protein localization in the eye. Single mutants of BBS subunits are viable and fertile. Reduced male fertility was observed only by the combined loss of BBS1 and BBS9. We found that the mutants retain gross motor coordination but show reduced positive phototaxis, while light-dependent modulation of locomotor activity remains detectable under the conditions examined. Tagged BBSome subunits localize to shared puncta in differentiating photoreceptors. Moreover, combined loss of BBSome alters the distribution of a rhodopsin reporter in adult photoreceptors, and leads to additional accumulation of the reporter in the cell-body region. Together, these findings indicate a nonciliary contribution of BBSome proteins to rhodopsin organization and visual function.

## Materials and Methods

### Drosophila stocks

Drosophila stocks were maintained on standard yeast-cornmeal-sucrose medium at 25 °C. CRISPR/Cas9 mutagenesis was used to generate deletion mutants for BBSome subunits. Two guide RNA targeting sequences targeting sequences close to the 5’ and 3’ ends of BBSome subunit coding sequences were tandemly cloned into pCFD5_w Drosophila guide RNA expressing vector as described in Port and Bullock, 2016. Guide RNA vectors were integrated into the genome either on attP40 or attP2 landing sites. Mutagenesis was performed by crossing males from these lines to nos-Cas9 virgins. Candidate lines were established from F2 generations, screened for deletions by PCR. Mutations breakpoints were then determined by Sanger sequencing.

Indels in the BBS18 coding sequence were generated by single gRNA strategy using TKO.GS01113 line from Bloomington Drosophila Stock Center (BDSC ID: 77279). Expression vectors for mRFP and mNeonGreen tagged contructs were generated by cloning BBSome subunit coding sequences into pUWRattB and pTiDNattB vectors by the Gateway method. Subsequent expression vectors were injected into phiC31; attp40 and phiC31; attP2 lines and transgenic stocks were established.

### Fertility tests

To test fertility, virgin females and males were collected and individually mated with two Oregon-R partners (ages 2–4 days). The crosses were kept at 25 °C for 6 d and the adults then removed. Numbers of eclosed progeny in each vial were recorded and statistically analyzed. Vials in which any adults died were not considered. We evaluated 15 crosses per genotype.

### Testing coordination

To quantitatively evaluate coordination, cohorts of 15 freshly eclosed flies were transferred into fresh vials with media and aged for 4 days. Immediately before the assay, flies were transferred without anesthesia into a clear empty testing vial. For the assay, vials were illuminated from above, flies were gently tapped down to the bottom of the vial, and then given 5 seconds to climb up the vial. Numbers of flies crossing the 5 cm mark were then recorded. The climbing assay was repeated three times for each cohort with similar results. Three independent cohorts of 15 flies were tested per genotype.

### Confocal microscopy

For fluorescent imaging, eye discs from wandering L3 larvae or pharenocephalic pupae were dissected in PBS and fixed in 4% formaldehyde for 20 min. Samples were than washed and mounted on microscope slides in Vectashield+DAPI. Samples were imaged using a Leica Stellaris 8 confocal laser scanning microscope with a 63× objective. Recorded images were opened in ImageJ (https://imagej.nih.gov/ij/), and colocalization was assessed by overlapping signals in merged color channels.

### Locomotor activity measurement

Daily pattern of locomotor activity and sleep were analyzed using the Drosophila Activity Monitoring System (DAMS; Trikinetics) as described in Skoczek et al, 2026.

### Cryosectioning of adult eyes

For cryosections, fly were decapitated and the heads were fixed in 4% paraformaldehyde for 4 h, followed by cryoprotection (2 × 10min in PBS, 1 × 10 min in 12.5% sucrose, and overnight in 25% sucrose). Samples were then mounted in Cryomatrix and frozen in liquid nitrogen. Sections of 20 μm thickness were prepared and mounted in Vectashield medium and imaged using a Zeiss laser-scanning microscope.

## Results

### BBSome subunit null mutants are viable and fertile

To investigate the role of BBS proteins in sensory functions, we first identified Drosophila homologs of the human proteins using FlyBase annotation and independently evaluated their homology. The targeted set comprised BBS1, BBS4, BBS5, BBS8, BBS9, and BBS18, together with the BBSome-associated GTPase BBS3/ARL6. This distinction is relevant when interpreting the phenotypes because BBS3 regulates the complex rather than forming part of its structural core (Jin et al, 2010). For brevity, we refer to this collection of BBSome-subunit and BBS3 mutants as BBS mutants. In accordance with the FlyBase BBSome group record, no homologs were identified for subunits BBS2 and BBS7.

To abolish gene function for Drosophila orthologues of BBsome subunits, we used CRISPR/Cas9 mutagenesis. Pairs of guide RNAs were used to target both 5’ and 3’ ends of the coding sequences to generate null mutants. The only exception was BBS18. Because this gene has a small coding region, we used a single guide RNA to generate a two-base-pair indel leading to a frame shift mutation. PCR analysis detected the deletion-containing fragments, and T7 endonuclease I digestion supported the presence of the BBS18 indel. Sequencing determined the mutation breakpoints (Figure 1). These alleles provided a panel of putative null mutants for subsequent phenotypic analysis.

**Fig 1.**
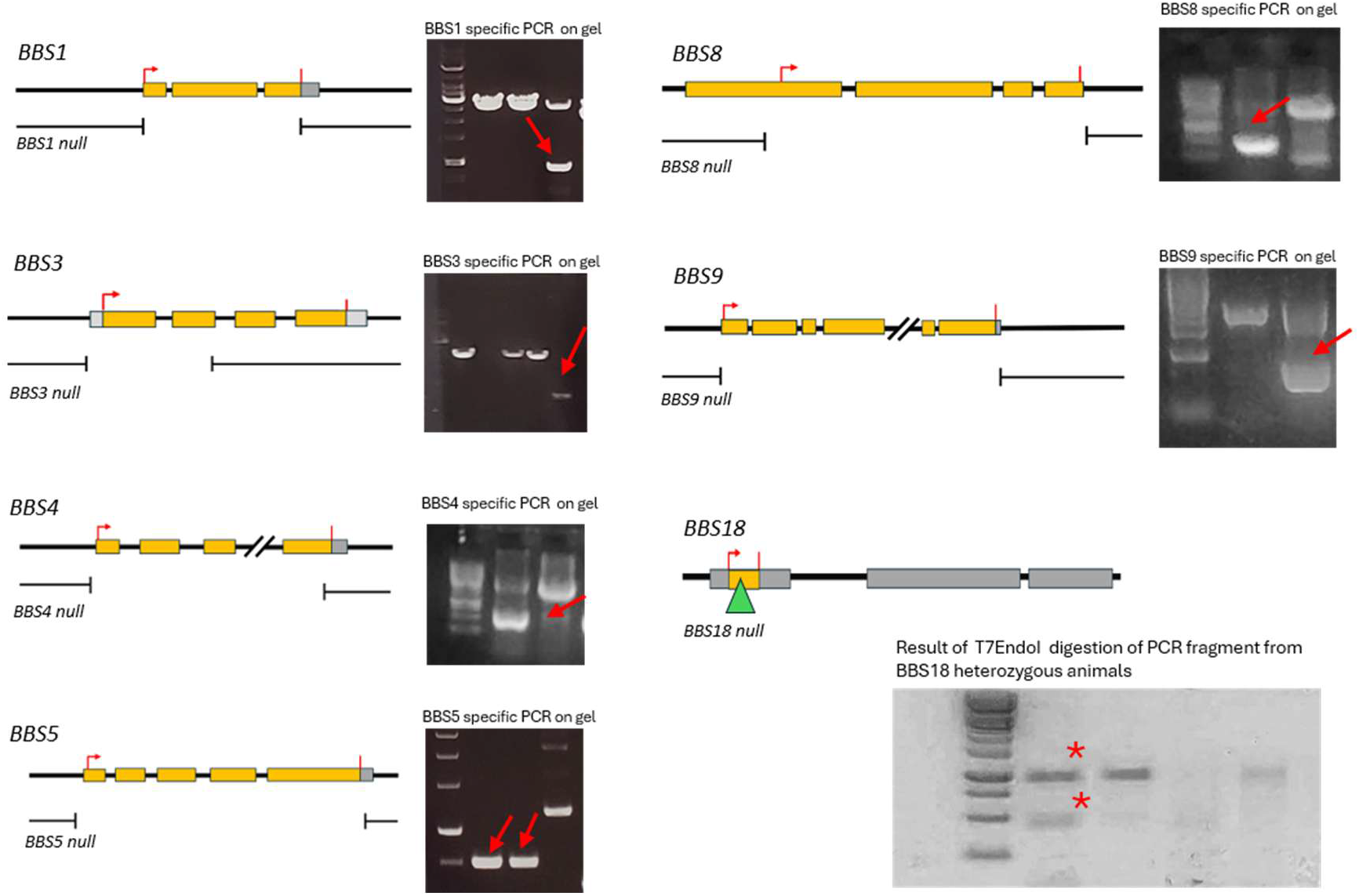
Molecular map of CRISPR/CasG generated null mutants. A structural map of each BBS gene is shown. The breakpoints of double-gRNA generated mutations are indicated under the map. One exception is BBS18, which was generated by a single gRNA induced indel (green triangle: the position of the 2bp indel). Alongside the genetic map, a representative agarose gel indicates the presence of mutant PCR fragments (red arrows: PCR fragment with deletion, red asterisk: T7EndoI digested PCR fragments with indel). Precise breakpoints the mutations were determined by Sanger sequencing.

Systematic preliminary characterization of homozygotes reveled, that all single mutants survived to adulthood and were fertile. Double mutant combinations between the WD40 and gamma-adaptin ear domain containing Drosophila BBSome subunits, BBS1 and BBS9 also resulted in complete viability (Figure 2A). Both female and male fertility were measured by mating mutant individuals to Oregon-R control line. Both sexes of each single mutant line produced substantial numbers of progeny, not significantly different from wild type controls (Figure 2B). In contrast, BBS1-BBS9 double-mutant males showed a substantial decrease in fathering progeny. Thus, the targeted BBS genes were individually dispensable for viability and fertility under, whereas combined loss of BBS1 and BBS9 revealed a strong reproductive phenotype.

**Fig 2.**
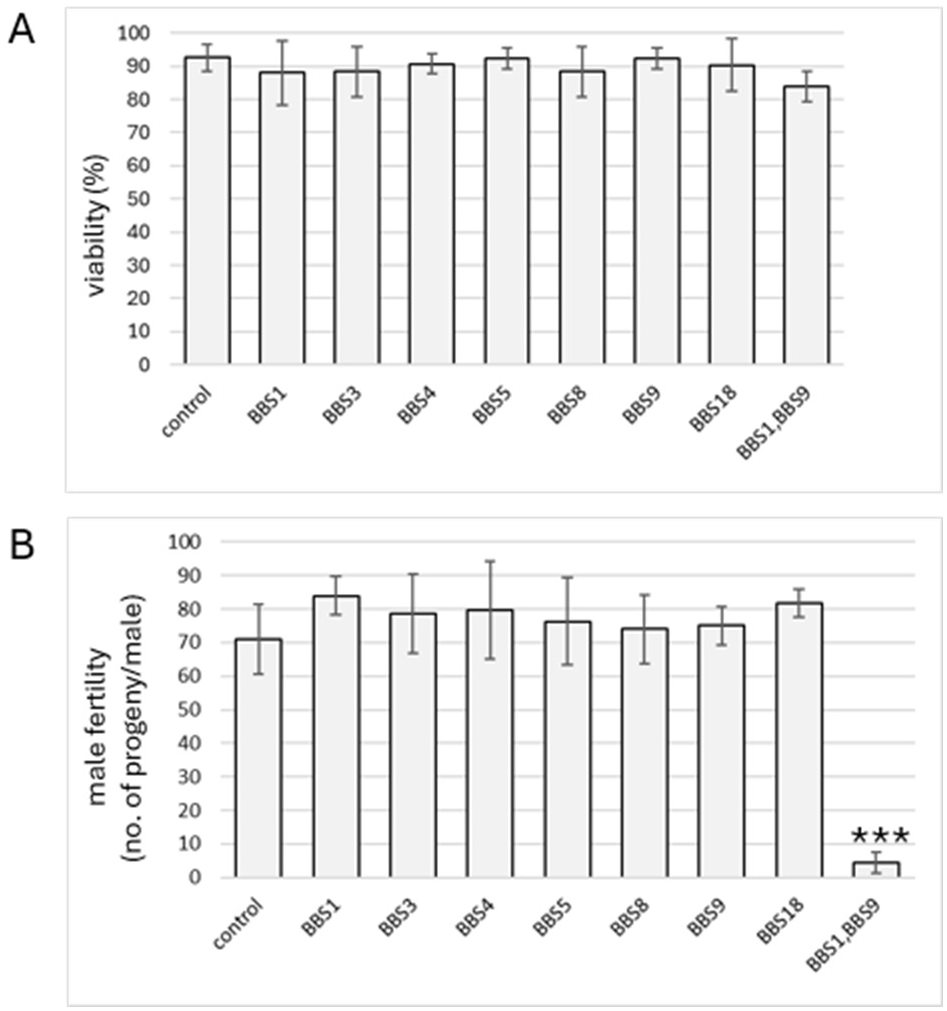
Viability and fertility of BBSome mutants. A) viability parameters – survival of L1 larvae to adulthood - of individual BBSome mutants are indicated. In each experiment, 200 L1 larvae were collected per genotype. Average and standard deviation of three independent biological replicates are indicated. B) Fertility of males – no. of eclosed progeny from each mutant males mated with 2 Oregon-R virgins – are indicated. In each experiment, the fertility of 10 males per genotype were tested. Average and standard deviation of three independent biological replicates are indicated. ***p< 0.001.

### BBsome mutants do not show functional ciliary abnormalities

Previous work identified BBS1, BBS4, and BBS8 among cilia-associated genes expressed in *Drosophila* mechanosensory and chemosensory organs (Avidor-Reiss et al, 2004). Because sensory cilia in chordotonal organs contribute to proprioception and coordinated movement, we first asked whether BBS loss produced a gross motor phenotype consistent with severe sensory dysfunction. We assessed climbing after flies were tapped to the bottom of a vial and compared the BBS mutants with animals subjected to ubiquitous knockdown of the transition-zone protein Cep290.

As expected upon severe ciliary disfunction, Cep290 RNAi flies were severely uncoordinated and largely remained near the bottom of the vial, whereas BBS mutants climbed readily comparable to control (Figure 3A). Quantification of the fraction of animals climbing 5 cm confirmed the marked impairment of the Cep290 RNAi control and the retention of climbing ability across the BBSome mutant panel, including BBS1-BBS9 double mutants (Figure 3B). In contrast to vertebrate models, these results argue against the essential functional contribution of BBSome in Drosophila sensory cilia.

**Fig 3.**
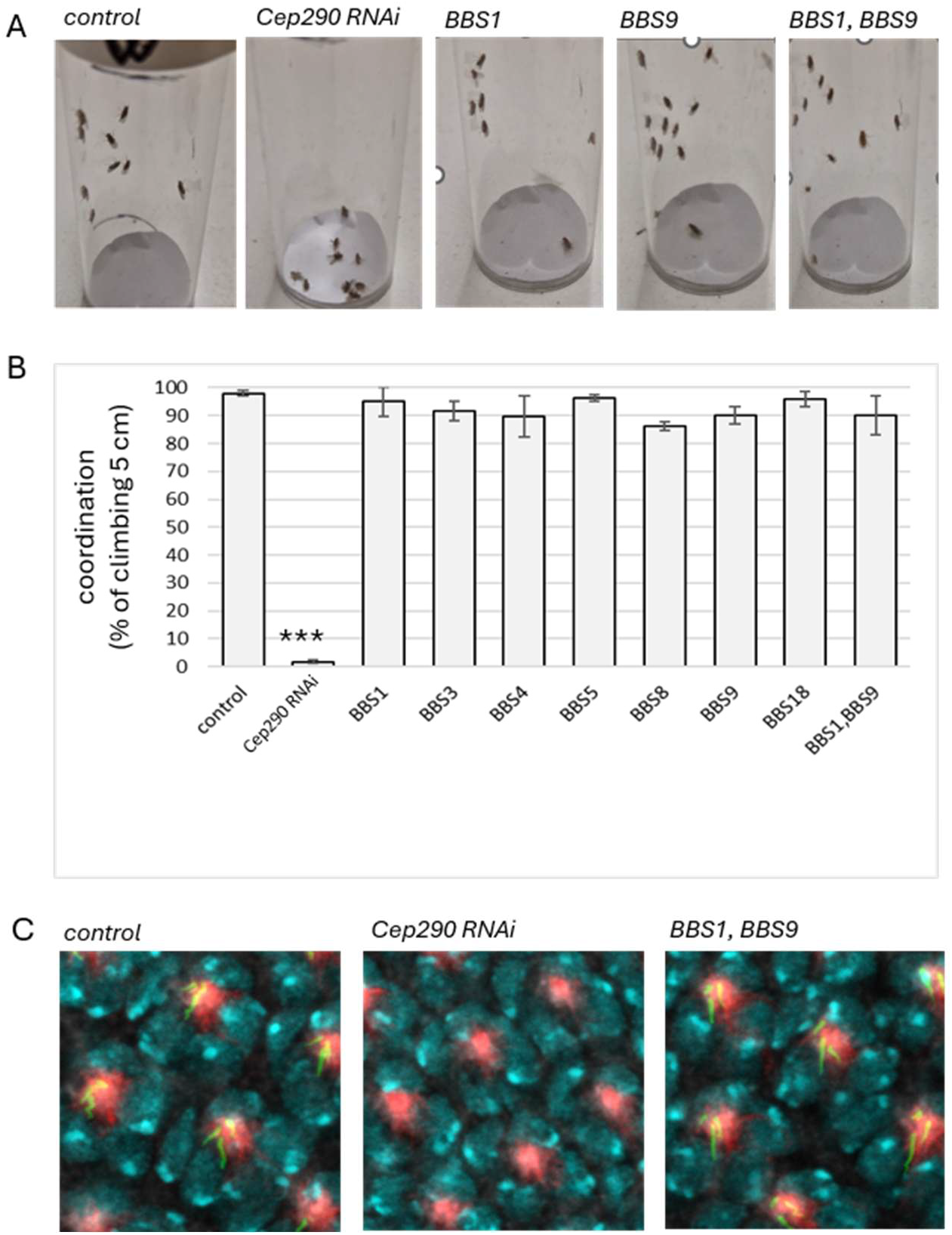
Lack of ciliary phenotype in BBsome mutants. **A)** Snapshots from climbing assays of Cep290 RNAi and BBS1 mutants. Images taken 2 seconds after taping down the vials. B) Ǫuantification of climbing assay of Cep290 RNAi and BBS null mutants. Data represent mean and standard deviation of three biological replicates. Each replicate consisted of vials containing 15 flies. *** p<0.001. C) Cilia in interommatidial mechanosensory bristles revealed by GFP-Rootletin (green) in wild type control, ubiquitously induced Cep290 RNAi and BBS1,BBS9 double null mutant pupal eye. Red, actin (Alexa568-Phalloidin), Cian, nuclei (DAPI).

To complement the analysis of coordination, we examined ciliary structures in the sensory preparations shown in Figure 3C. Rootletin organizes the ciliary rootlet, a structure associated with the basal apparatus of sensory cilia and is therefore informative about ciliogenesis (Chen et al, 2015). Controls displayed discrete elongated GFP-Rootletin structures. Corresponding structures were not readily detectable in the Cep290 RNAi preparation, whereas they remained intact in BBSome mutants, including the BBS1-BBS9 double mutant (Figure 3C). This evidence combined with the functional climbing assays suggest that Drosophila sensory cilia retain rootlet-associated ciliary organization in absence of the BBSome.

### BBsome mutants exhibit visual defects

The connection between BBSome function and vision in vertebrates is explained by the role of this protein complex in intraciliary transport within the connecting cilium of photoreceptor cells. The fact that *Drosophila* photoreceptors lack a connecting cilium or any similar structure, led us to ask whether the BBSome might still have some role in visual processes.

An initial indication of altered visual behavior was a weaker startle response by BBS mutants to lights being switched off (Supplementary Video 1). We therefore tested the preferential movement of flies toward light using a multiple-choice maze phototaxis assay adapted from Bosch and colleagues (Bosch et al, 2015). Flies entered through the lower opening and moved through successive choices to one of nine exit positions. Exit positions were assigned scores from −4 to +4, with increasingly positive scores corresponding to exits closer to the light source (Figure 4A).

**Fig 4.**
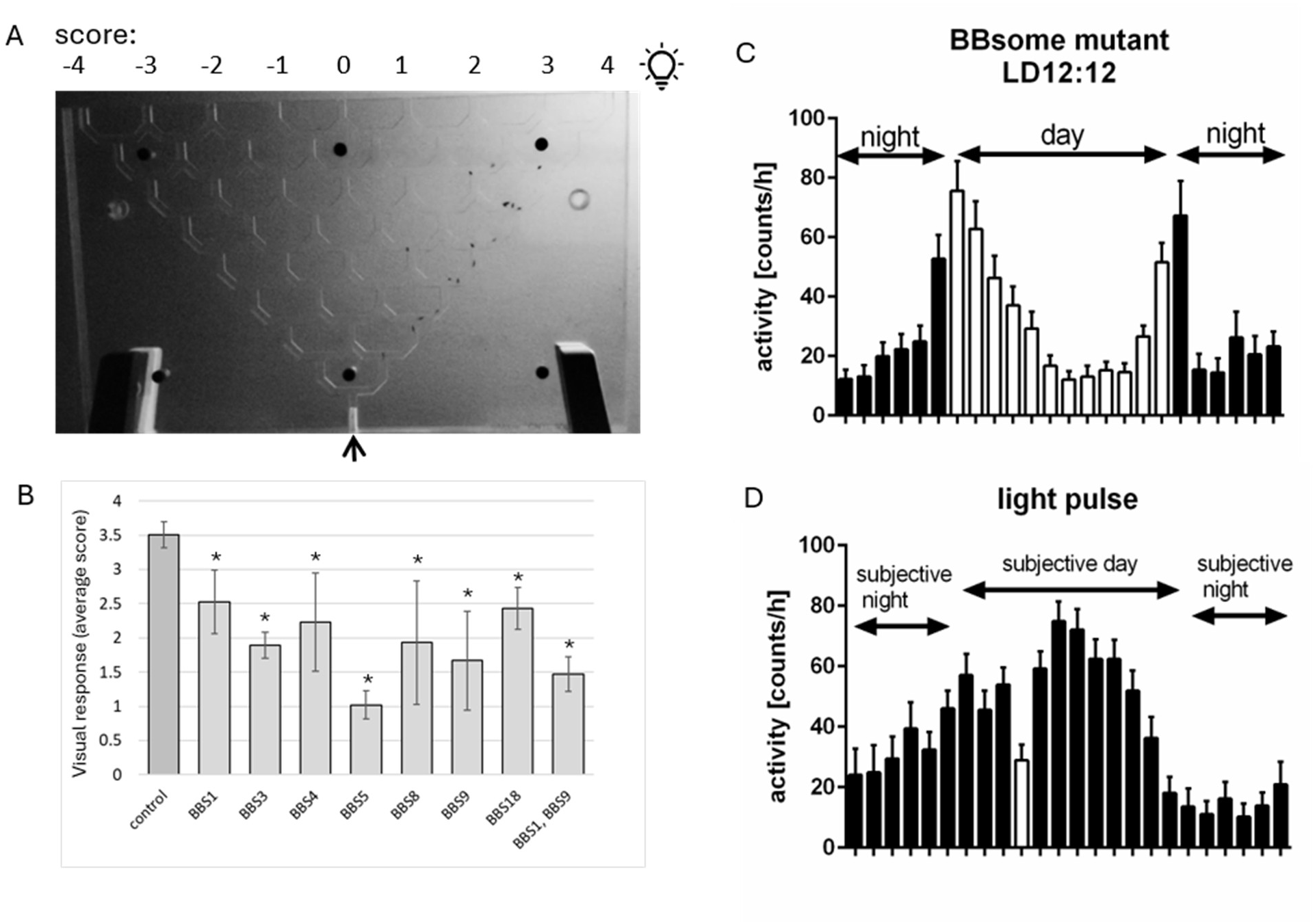
Visual abnormalities in BBSome mutants. A) The set up of Drosophila phototaxis assay modified after Bosch et al, 2015 to quantitatively test visual response. Flies entering the maze (bottom hole, arrow) exit one of the nine holes on top. Positive scores indicate the proximity to the light source (lightbulb icon). B) Average visual response values of 30 flies from each genotype. Data represent mean and standard deviation. C) Locomotor activity of BBSome mutant flies under LD12:12 conditions (12 h light:12 h dark). Flies exhibited a typical bimodal activity pattern, with morning and evening activity peaks occurring at ZT1 (1 h after lights on) and ZT13 (1 h after lights off), respectively. D) BBSome mutant flies retain responsiveness to light. After 3 days in constant darkness (DD), a 1 h light pulse administered during the subjective day induced a transient decrease in locomotor activity. This reduction was observed only during light exposure, with activity levels returning to baseline immediately after the light pulse.

In this assay, control flies showed a strong preference for the illuminated side of the maze. Every BBS mutant genotype tested had a lower mean phototaxis score, including the BBS1 BBS9 double mutant (Figure 4B). The mean scores remained positive, indicating that the response to light was reduced rather than completely lost. Because the same mutants retained normal climbing ability, the phototaxis phenotype is unlikely to be explained by the locomotor incapacity. The assay nevertheless indicates a significantly decreased behavioral preference towards light.

We next examined locomotor activity in relation to the lighting regime. Flies entrained to a 12 h light:12 h dark cycle exhibited a bimodal locomotor activity pattern, with activity peaks at ZT1 and ZT13, where ZT0 denotes lights on and ZT12 denotes lights off (Figure 4C). In the mutant condition examined, activity was elevated during the first half of the light phase relative to the middle of the day. The persistence of a structured daily activity profile showed that the phototaxis defect was compatible with organized locomotor activity under light–dark conditions.

Following transfer to constant darkness (DD), flies exhibited free-running locomotor rhythms. Interestingly, the light pulse did not elicit a startle response, as no transient increase in locomotor activity was observed immediately after light onset. Instead, activity decreased during the pulse. This finding suggests that BBSome mutants retain light responsiveness, but the response is predominantly inhibitory, consistent with a negative masking effect rather than an acute locomotor activation induced by light. Alternatively, light exposure could induce a transient freezing-like behavioral response or promote sleep through activation of homeostatic sleep mechanisms.

Collectively, these observations indicate that BBSome loss impairs aspects of visually guided behavior without eliminating all behavioral responses to light.

### BBSome subunits localize to shared puncta in differentiating photoreceptors

The altered visual responses in BBSome mutants led us to ask whether the localization of BBSome subunits was consistent with a role in the developing retina. During the third larval instar, photoreceptor differentiation proceeds posterior to the advancing morphogenetic furrow in the eye imaginal disc. Fluorescently tagged BBSome proteins were detectable in this differentiating region, where they formed discrete puncta within the developing photoreceptor clusters (Figure 5A). To test whether different subunits are incorporated into the same structures, we coexpressed mNeonGreen-tagged BBS1 and RFP-tagged BBS4. Their signals overlapped in discrete puncta in differentiating photoreceptors (Figure 5B), consistent with recruitment of multiple BBSome proteins to a shared subcellular compartment.

**Fig 5.**
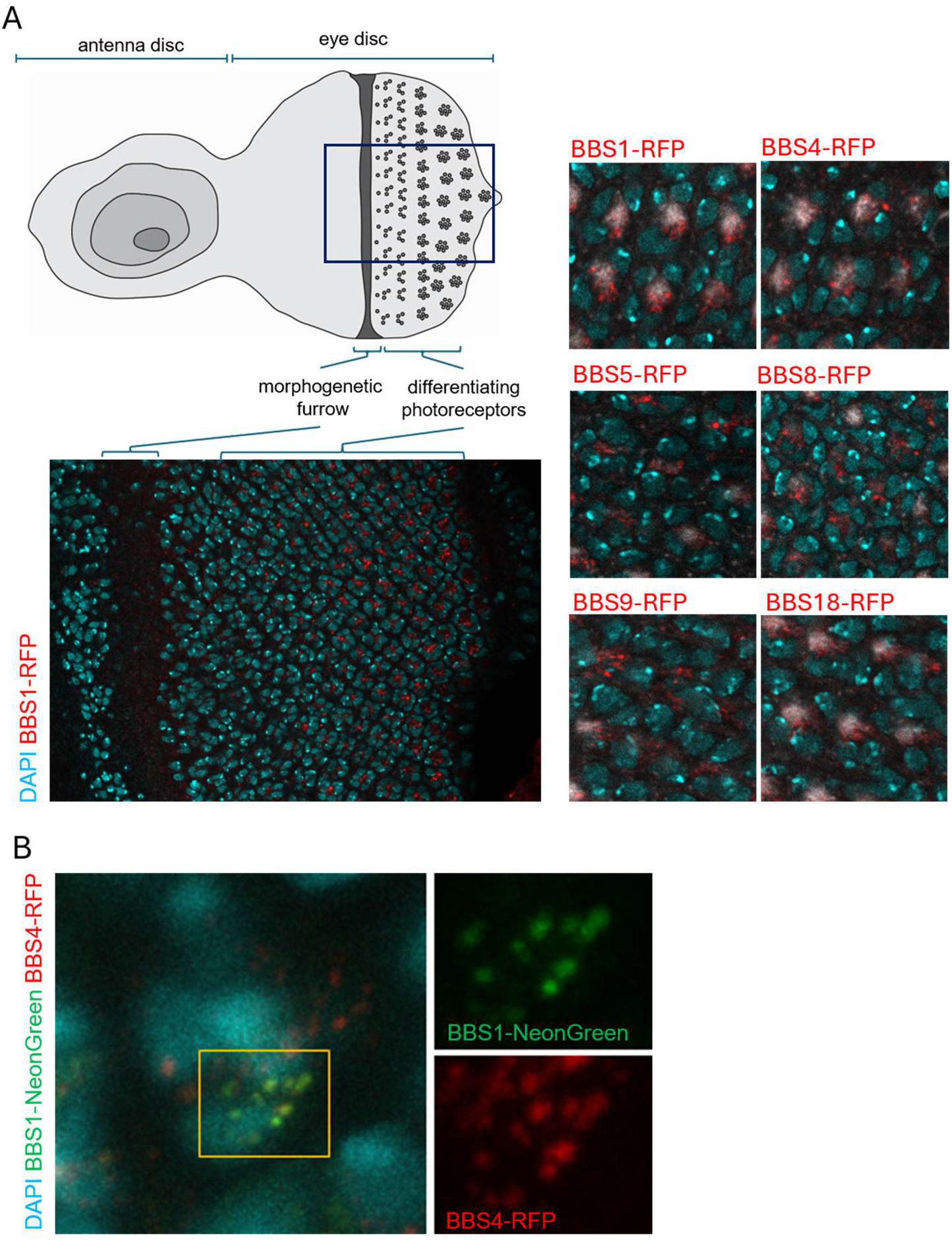
BBSome subunits are expressed and localize to differentiating photoreceptor cells. A) RFP tagged BBS subunit expresses after the morphogenetic furrow and form dot-like structures in differentiating photoreceptor cells of wandering L3 larva. Higher magnification panels indicate the localization of each tested subunit. B) Colocalization of RFP-tagged BBS4 and mNeonGreen-tagged BBS1 subunits indicating the assembly of the complex in differentiating photoreceptors.

Biochemical association provided complementary support for complex formation in eye tissue. Pulldowns using GMR-GAL4-driven UAS-mNeonGreen-tagged BBS1, BBS8, or BBS9 recovered other BBSome subunits (Supplementary Table 1). Together with the overlapping localization of BBS1 and BBS4, these results support the association of BBSome proteins in the developing eye. The puncta are therefore plausible sites of BBSome-associated activity, although their precise identity and the stoichiometry of subunits remain unresolved in this tissue.

### Loss of BBSome alters rhodopsin reporter distribution

To determine whether BBSome mutations would affect the mechanism localizing photoreceptor, we followed the distribution of Rhodopsin-1, the major photosensitive molecule in photoreceptor cells, encoded by the *ninaE*. To this end, we compared control and BBS1-BBS9 double-mutant photoreceptors in Drosophila eyes expressing a *ninaE-GFP* transgene under its own promoter. In cryosections of control eyes, ninaE-GFP fluorescence was concentrated in the elongated rhabdomeres, with relatively little signal in photoreceptor cell bodies (Figure 6). However, at matched acquisition setting, double-mutant eyes showed an increased rhabdomeric fluorescence with the addition of conspicuous GFP-positive puncta in the cell-body region (arrow, Figure 6). This redistribution is consistent with defective rhodopsin trafficking or turnover. Importantly, as Drosophila retinal photoreceptors lack cilia, the altered intracellular distribution of Rhodopsin-1 in BBS mutants indicates a non-ciliary function of BBSome in insect photoreceptors.

**Fig 6.**
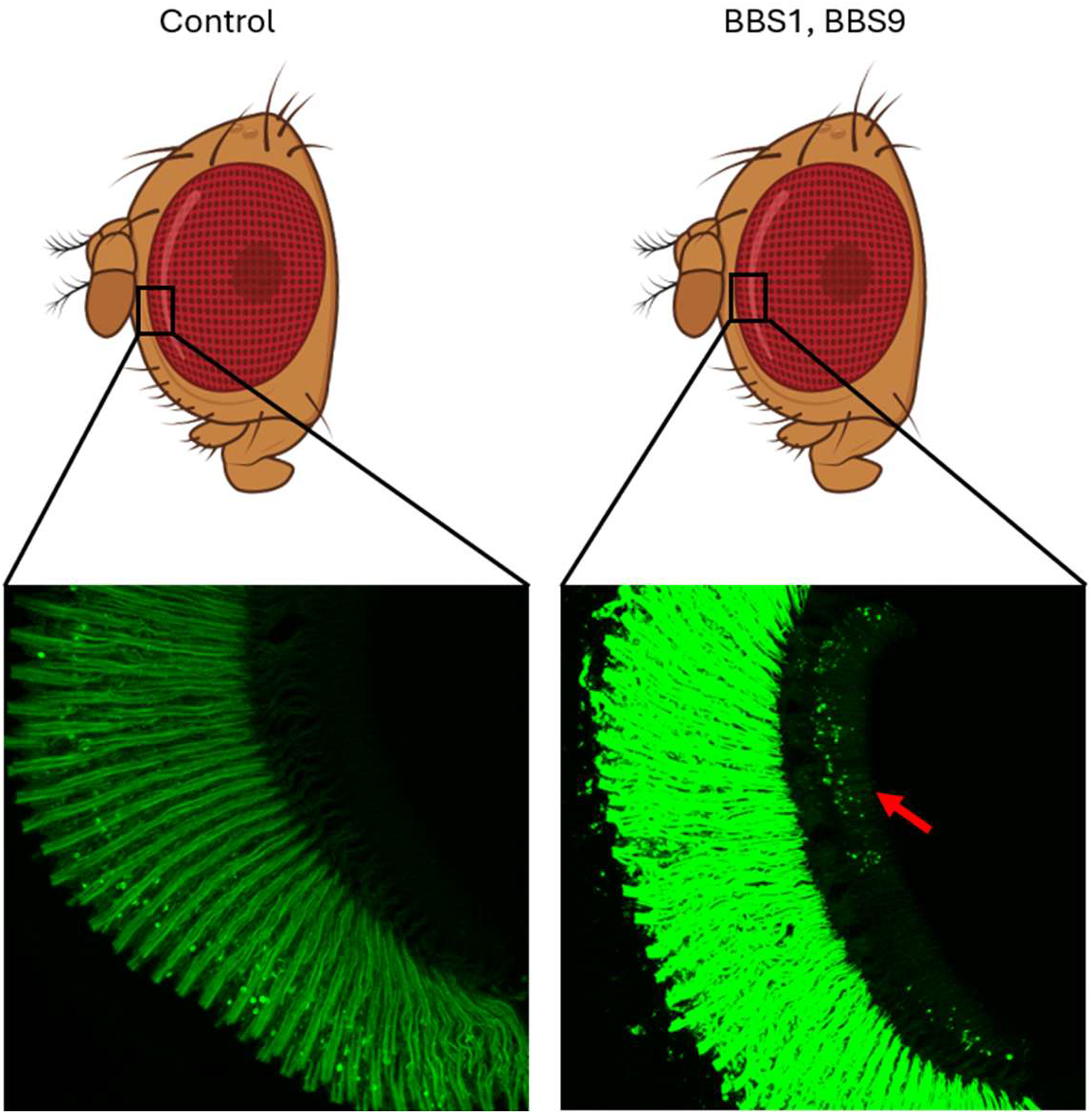
Accumulation of ninaE in rhabdomers and photoreceptor cell bodies. Cryosections of control and BBS1-BBS9 double mutant adult eyes expressing transgenic ninaE-GFP from its own promoter. Elongated rhabdomere stripes are visible. Arrow indicate accumulation of ninaE positive puncta in photoreceptor cell bodies.

## Discussion

Retinal degeneration in Bardet–Biedl syndrome has generally been considered a consequence of defective trafficking in the connecting cilium of vertebrate photoreceptors. Studies in mouse models suggest a requirement for BBSome function in outer-segment formation, maintenance and protein composition of photoreceptor cells (Datta et al., 2015; Hsu et al., 2017). Whether the complex also contributes to visual function independently of the ciliary compartment has remained less clear. The Drosophila retina provides an opportunity to address this question because its photoreceptors concentrate rhodopsin in rhabdomere microvilli and there is no connecting cilium. Here we show that mutations affecting BBSome subunits, or the associated regulator BBS3, reduce visually guided behavior despite the retention of gross motor coordination. We also found that BBSome subunits are expressed in differentiating photoreceptors to accumulate in distinct punctate. In wild type photoreceptors, the Rhodopsin1-GFP reporter has a uniform distribution in rhabdomeres and there is virtually no signal in cell bodies. However, In BBSome mutant adults, the rhodopsin reporter accumulates extensively in rhabdomeres and in addition, forms punctate structures in photoreceptor cell bodies. Together, these observations support a nonciliary contribution of BBSome proteins to retinal function.

They suggest that the relationship between the BBSome and vision is not confined to transport through a connecting cilium, but may also involve intracellular processes that maintain the distribution of photosensitive membrane proteins. These observations connect a behavioral phenotype to an abnormality of Rhodopsin distribution within photoreceptors that is independent of ciliary trafficking.

The established molecular properties of the BBSome provide a basis for such a function associated with cilia in the mammalian retina. The complex assembles a membrane-associated coat under the control of ARL6/BBS3 and recognizes membrane protein cargo, while BBSome trains mediate the removal of activated receptors from cilia (Jin et al., 2010; Ye et al., 2018). Our localization and pulldown results suggest that conserved BBSome components also associate in the nonciliated photoreceptor cells. The colocalization of BBS1 and BBS4 in punctae is supported by the recovery of additional subunits with BBS1, BBS8 and BBS9 baits in pull downs. These findings are consistent with a complex acting at intracellular sorting sites, although the identity of those sites and their association with rhodopsin remains to be established. Notably, we could not identify any Drosophila orthologue for BBS2 and BBS7. The functional complexes of the fly eye therefore do not necessarily have the exact composition of the mammalian BBSome. It is reasonable to suppose that the conserved components participate in membrane protein sorting in this tissue, but colocalization and biochemical association alone do not establish the stoichiometry of the complex or a direct interaction with ninaE/Rh1.

The altered distribution of the rhodopsin reporter does, however, provide a cellular indication of disrupted membrane protein homeostasis. Rhodopsin 1 normally reaches the rhabdomere through polarized transport and subsequently undergoes light-dependent endocytosis and recycling. Rab11 is required for its post-Golgi delivery, whereas the retromer retrieves endocytosed Rhodopsin 1 and protects photoreceptors from the consequences of abnormal endolysosomal processing (Satoh et al., 2005; Wang et al., 2014). In our BBSome mutants, the additional punctae in cell-bodies are accompanied by stronger rhabdomeric reporter fluorescence under matched acquisition settings. This combination suggests altered partitioning or turnover of the reporter, rather than its simple absence from the photosensitive membrane. However, our images do not establish whether the puncta contain newly synthesized or internalized Rhodopsin 1, or material destined for degradation. Nor does reporter fluorescence alone establish an increase in total endogenous rhodopsin abundance. We therefore favour a model in which BBSome loss disturbs the balance between rhabdomeric and intracellular Rhodopsin pools, with defective recycling representing the most plausible explanation. Future characterization of the behaviour of Rhodopsin 1 after controlled illumination would distinguish this possibility from defective biosynthetic transport, altered endocytosis or impaired degradation. Such experiments would also determine whether the puncta occupied by tagged BBSome subunits in the developing retina are related to the compartments that accumulate rhodopsin in adult mutants. Our striking finding is that the developmental localization of BBSome components, and the consequences of their genetic elimination upon the distribution of the Rhodopsin reporter and ability of flies to respond to light strongly support a retinal role for BBSome in an unexpected nonciliary context

## Aknowledgement

Weare grateful to Dr. Andres Collazo and Dr. Giada Spigolon of the Biological Imaging Facility at Caltech for their support and to Caltech Beckman Institute Cryo-EM Facility for the use of the Tecnai electron microscope. We are thankful to Paula Coelho, Agota Nagy, and Pallavi Panda for many discussions, and to Orsolya Dózsa for occasional technical help provided. Pilot studies for this work were carried out with grant support from the NIH (Grants R01NS119614 and R01GM143699) and from the Caltech Center for Evolutionary Science to David M Glover. The body of this work was supported by PN-IV-P2-2_1-TE-2023-0123 grant from UEFISCDI office of Romanian Ministry of Education and Research to Levente Kovacs. Milena Damulewicz was supported by NCN UMO-2022/47/B/NZ3/00250 grant from Poland’s National Science Centre.

## References

Avidor-Reiss, T., Maer, A. M., Koundakjian, E., Polyanovsky, A., Keil, T., Subramaniam, S., & Zuker, C. S. (2004). Decoding cilia function: defining specialized genes required for compartmentalized cilia biogenesis. Cell, 117(4), 527–539. 10.1016/s0092-8674(04)00412-x

Beales, P. L., Elcioglu, N., Woolf, A. S., Parker, D., & Flinter, F. A. (1999). New criteria for improved diagnosis of Bardet-Biedl syndrome: results of a population survey. Journal of medical genetics, 36(6), 437–446.

Bosch, D. S., van Swinderen, B., & Millard, S. S. (2015). Dscam2 affects visual perception in Drosophila melanogaster. Frontiers in behavioral neuroscience, 9, 149. 10.3389/fnbeh.2015.00149

Cassioli, C., Onnis, A., Finetti, F., Capitani, N., Brunetti, J., Compeer, E. B., Niederlova, V., Stepanek, O., Dustin, M. L., & Baldari, C. T. (2021). The Bardet-Biedl syndrome complex component BBS1 controls T cell polarity during immune synapse assembly. Journal of cell science, 134(16), jcs258462. 10.1242/jcs.258462

Chen, J. V., Kao, L. R., Jana, S. C., Sivan-Loukianova, E., Mendonça, S., Cabrera, O. A., Singh, P., Cabernard, C., Eberl, D. F., Bettencourt-Dias, M., & Megraw, T. L. (2015). Rootletin organizes the ciliary rootlet to achieve neuron sensory function in Drosophila. The Journal of cell biology, 211(2), 435–453. 10.1083/jcb.201502032

Datta, P., Allamargot, C., Hudson, J. S., Andersen, E. K., Bhattarai, S., Drack, A. V., Sheffield, V. C., & Seo, S. (2015). Accumulation of non-outer segment proteins in the outer segment underlies photoreceptor degeneration in Bardet-Biedl syndrome. Proceedings of the National Academy of Sciences of the United States of America, 112(32), E4400–E4409. 10.1073/pnas.1510111112

FlyBase. BBSOME gene group report FBgg0000094. Accessed November 8, 2026. https://flybase.org/reports/FBgg0000094.html

Guo, D. F., Cui, H., Zhang, Q., Morgan, D. A., Thedens, D. R., Nishimura, D., Grobe, J. L., Sheffield, V. C., & Rahmouni, K. (2016). The BBSome Controls Energy Homeostasis by Mediating the Transport of the Leptin Receptor to the Plasma Membrane. PLoS genetics, 12(2), e1005890. 10.1371/journal.pgen.1005890

Hsu, Y., Garrison, J. E., Kim, G., Schmitz, A. R., Searby, C. C., Zhang, Q., Datta, P., Nishimura, D. Y., Seo, S., & Sheffield, V. C. (2017). BBSome function is required for both the morphogenesis and maintenance of the photoreceptor outer segment. PLoS genetics, 13(10), e1007057. 10.1371/journal.pgen.1007057

Jin, H., White, S. R., Shida, T., Schulz, S., Aguiar, M., Gygi, S. P., Bazan, J. F., & Nachury, M. V. (2010). The conserved Bardet-Biedl syndrome proteins assemble a coat that traffics membrane proteins to cilia. Cell, 141(7), 1208–1219. 10.1016/j.cell.2010.05.015

Loktev, A. V., Zhang, Q., Beck, J. S., Searby, C. C., Scheetz, T. E., Bazan, J. F., Slusarski, D. C., Sheffield, V. C., Jackson, P. K., & Nachury, M. V. (2008). A BBSome subunit links ciliogenesis, microtubule stability, and acetylation. Developmental cell, 15(6), 854–865. 10.1016/j.devcel.2008.11.001

Nachury, M. V., Loktev, A. V., Zhang, Q., Westlake, C. J., Peränen, J., Merdes, A., Slusarski, D. C., Scheller, R. H., Bazan, J. F., Sheffield, V. C., & Jackson, P. K. (2007). A core complex of BBS proteins cooperates with the GTPase Rab8 to promote ciliary membrane biogenesis. Cell, 129(6), 1201–1213. 10.1016/j.cell.2007.03.053

Satoh, A. K., O’Tousa, J. E., Ozaki, K., & Ready, D. F. (2005). Rab11 mediates post-Golgi trafficking of rhodopsin to the photosensitive apical membrane of Drosophila photoreceptors. Development (Cambridge, England), 132(7), 1487–1497. 10.1242/dev.01704

Skoczek, K., Mitka, M., Pyza, E., & Damulewicz, M. (2026). When nights are no longer dark: Aging and light pollution reshape visual system rhythms. npj Aging. 10.1038/s41514-026-00516-0

Wang, S., Tan, K. L., Agosto, M. A., Xiong, B., Yamamoto, S., Sandoval, H., Jaiswal, M., Bayat, V., Zhang, K., Charng, W. L., David, G., Duraine, L., Venkatachalam, K., Wensel, T. G., & Bellen, H. J. (2014). The retromer complex is required for rhodopsin recycling and its loss leads to photoreceptor degeneration. PLoS biology, 12(4), e1001847. 10.1371/journal.pbio.1001847

Ye, F., Nager, A. R., & Nachury, M. V. (2018). BBSome trains remove activated GPCRs from cilia by enabling passage through the transition zone. The Journal of cell biology, 217(5), 1847–1868. 10.1083/jcb.201709041

Port, F., & Bullock, S. L. (2016). Augmenting CRISPR applications in Drosophila with tRNA-flanked sgRNAs. Nature methods, 13(10), 852–854. 10.1038/nmeth.3972

